# Transfer learning framework for cell segmentation with incorporation of geometric features

**DOI:** 10.1101/2021.02.28.433289

**Authors:** Yinuo Jin, Alexandre Toberoff, Elham Azizi

**Affiliations:** Department of Computer Science, Columbia University, New York, NY, USA; Department of Biomedical Engineering and Irving Institute for Cancer Dynamics, Columbia University, New York, NY, USA

## Abstract

With recent advances in multiplexed imaging and spatial transcriptomic and proteomic technologies, cell segmentation is becoming a crucial step in biomedical image analysis. In recent years, Fully Convolutional Networks (FCN) have achieved great success in nuclei segmentation in *in vitro* imaging. Nevertheless, it remains challenging to perform similar tasks on *in situ* tissue images with more cluttered cells of diverse shapes. To address this issue, we propose a novel transfer learning, cell segmentation framework incorporating shape-aware features in a deep learning model, with multi-level watershed and morphological post-processing steps. Our results show that incorporation of geometric features improves generalizability to segmenting cells in *in situ* tissue images, using solely *in vitro* images as training data.

## 1 Introduction

Studying the diversity and complex organization of cell types in tissues provides insight into their various biological functions, maintenance and transformation. Single-cell technologies are powerful in revealing the heterogeneity of cell types and cell states. However, they rely on tissue dissociation and are thus agnostic to the spatial context and intercellular interactions. More recent multiplexed imaging technologies with fluorescence *in situ* hybridization [9] provide the advantage of mapping the spatial context of cells in addition to profiling gene expression of individual cells. Emerging single cell proteomic technologies [9, 5] also allow for the direct study of protein expression levels.

With the rapid growth of these spatial imaging technologies, there is a strong demand for computational tools for analyzing this data. One crucial step prior to most downstream analysis is performing high-quality image segmentation. By successfully isolating each discrete cell in an image, we can perform other analyses such as quantifying the expression of each gene or protein in individual cells, which enables the characterization of cell types (e.g. by clustering cells based on expression profile) or cell state transitions (e.g. by trajectory inference) while accounting for the spatial organization and context of cells.

Despite recent success on specific cell imaging datasets [13, 6], Fully Convolutional Networks (FCN) models are unable to generalize across cell, culture, and tissue types; most success so far has been observed on symmetrical, well-separated *in vitro* nuclei stained cells [4]. Thus, developing pipelines for segmentation of *in situ* images which contain valuable biological information in the context of the tissue is an urgent, unmet need. Moreover, the scarcity of comprehensive annotated training datasets on tissue images, makes the generalizability of methods from the *in vitro* domain to other domains more essential.

### 1.1 Related Works

The U-Net architecture [13] is recognized as a popular and representative FCN for image segmentation in biomedical applications and beyond. However, to perform high-quality instance segmentation at the single cell resolution, especially for *in situ* tissue images, the base U-Net struggles with separating adjacent, clustered cells, leading to multiple methods that incorporate shape-aware priors during training or post-processing. Guerrero-Pena et al. [6], for example, designed a Shape-Aware Weight map (SAW), assigning higher weights to pixels corresponding to borders of attaching cells and lower weights to background regions in the cross-entropy loss calculation.

Traditionally, the Watershed algorithm, based on the concept of flooding from local minima to edges [2, 11], and accurate prediction of membrane markers at local minima is used to separate attached cells. Al-Kofahi et al. [1] expanded upon this by preceding the water-shedding with a deep learning output with *h*-minima and multiple rounds of Otsu-thresholding to separate connected cells in the prediction. Other approaches have attempted to learn the watershed energy levels in neural networks by performing distance transform to the ground-truth masks [3, 10].

Nevertheless, these methods have been either solely applied to certain image datasets (e.g. cluttered T-cells, Cityscapes), suffer from class imbalance or over-fitting issues, or are inflexible due to significant feature engineering. To address these issues, we propose a new approach that utilizes and adapts these ideas to incorporate geometric features as priors, while transferring knowledge attained from *in vitro* nuclei segmentation to the domain of *in situ* segmentation. This generalizability is important as there are many annotated *in vitro* nuclei segmentation training datasets available while *in situ* annotated data are rare, requiring time-consuming, expert manual segmentation.

## 2 Methods

### 2.1 Model Architecture

Our method utilizes a Feature Pyramid Networks (FPN) architecture with residual blocks (Figure 1)[12]. It maintains the contraction path from the U-Net architecture with major modifications to the extraction path. In every double-convolutional layer within the contraction path, a shortcut connection is added from the input to the output [7]. While U-Net-like architectures only make predictions at the last layer, the FPN utilizes feature maps from all resolution levels along the extraction path, which directly contribute to the output predictions.

**Figure 1:**
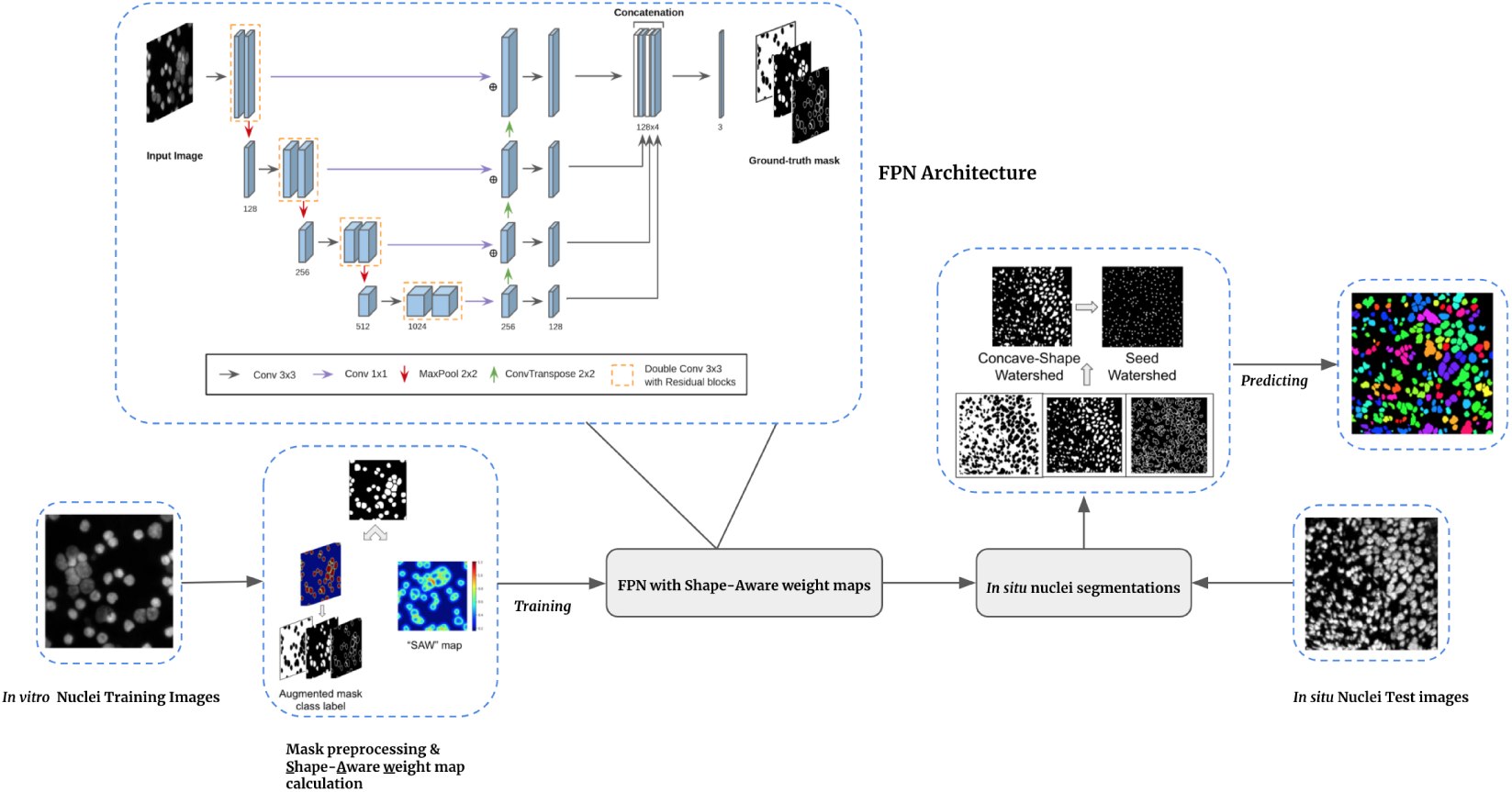
Overview of transfer learning approach with shape-aware priors in a Feature Pyramid Network and multi-level watershed post-processing

### 2.2 Preprocessing & Shape-Aware Weight Maps

We expanded upon the typical label augmentation for attaching cell borders [6] by assigning both the cell borders and contour pixels as a third label, besides background and nuclei. This modification combats class imbalance arising from border pixel sparsity in the *in vitro* training set, allowing our approach to generalize to various cell types.

We also modified the calculation of Shape-Aware Weight maps (SAW), which encodes geometric properties of cells in loss functions during training as following. For each pixel, the original equation of SAW Guerrero-Pena et al. [6] defines the weights as the sum of class imbalance weight and a “complexity” weight, which focuses on cell borders and narrow, irregular cell shapes:

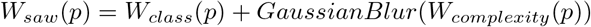

where *W*_*complexity*_ applies distance transformations as in Guerrero-Pena et al. [6] to each individual mask and its concave complement to address higher weights at smaller or concave cell regions. Gaussian blur (smoothing) is applied to assign relatively high weights to pixels close to those “complex” pixels. *L* ={0, 1, 2} denotes the set of all possible ground-truth labels, with integer labels from 0 to 2 representing background, foreground, and contour & attaching border pixels respectively. *g*_*i*_ ∈ *L* represents class labels and *W*_*class*_ assigns the class weight inversely proportional to the total counts of each class labels *ω*_*i*_ = 1*/*|*g*_*i*_|. However, for training sparse images with few cells, such class weight assignment would generate extremely low weights for background pixels, which may encourage the model to under-optimize at certain backgr<u>oun</u>d regions. Therefore, for each cI;lass label *g*_*i*_ ∈ *L*, we reassigned the class weights as 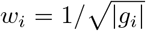 followed with normalization (∑_*i*_ *ω*_*i*_ = 1) to compensate for this problem.

For computational efficiency, both the label augmentation and the SAW maps for each input mask were calculated before the training (Figure 1). The pre-trained FPN model based on SAW with cross-entropy loss was then applied to a new domain involving *in situ* images with nuclei markers for testing.

### 2.3 Multi-level Watershed Post-processing

Despite the increase in accuracy over U-Net from the aforementioned model architecture with the incorporation of SAW, *in situ* intact tissue images with cluttered cells, especially when they are visually indistinguishable, are extremely difficult to accurately segment. We thus propose a fully automated multi-level watershed post-processing method (Algorithm 1) to detach connected cells in our FCN + SAW output. Given an arbitrary prediction map 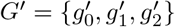 with three channel predictions for background (0), foreground (1), and cell boundary & attaching borders (2), we first select the predicted foreground regions, denoted as 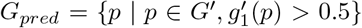. We then calculate the solidity^3^, which reflects the convexity level of each individual mask. For all masks with solidity higher than the cutoff parameter *θ* (i.e. “convex-like” masks), we measure the average area *a*_*avg*_ and diameter *d*_*avg*_ representing average cell size. All the other masks will be passed to the first-round watershed if their area exceed *a*_*avg*_. We set the closest possible distance between any two local minima in the watershed as *d*_*avg*_, the diameter learned from all convex-like masks. Such restrictions help avoid over-segmentation with false-positive local minima.

After the first-round watershed, we set the centroid of all candidate individual masks as the markers for the second-round watershed. The topography for watershed is set as the inverse distance transform^4^ from *G*_*pred*_, and the candidate regions for watershed are set as 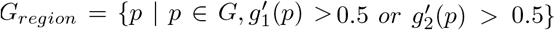 for all possible foreground, border or contour pixels. The second-round watershed recovers the maximized area of each individual nuclei mask and detects non-convex (e.g. bean-shaped concave cells).

Note that all the parameters in our multi-level watershed post-processing pipeline can be learned. *a*_*avg*_ and *d*_*avg*_ are estimated from each individual prediction maps during the first-round watershed. *θ* is calculated by the average solidity in the training set. In the Data Science Bowl training set[4], the average solidity is 93.90 so we roughly set *θ* = 0.9.

#### Algorithm 1

Multi-level Watershed Post-processing

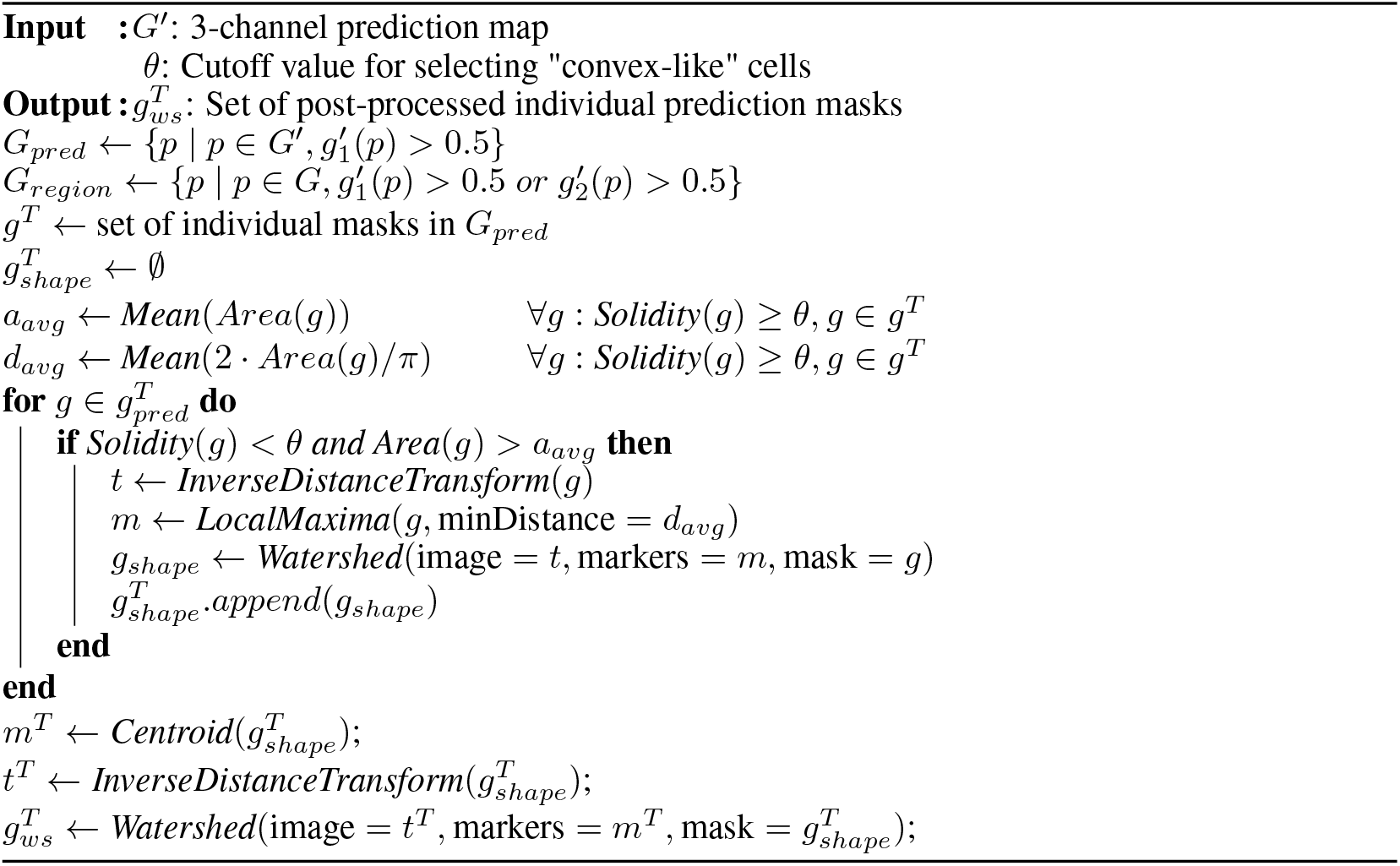

## 3 Results

### 3.1 Dataset

We gathered nuclei images from multiple sources for *in vitro* and *in situ* cells. *In vitro* cell images are downloaded from the 2018 Data Science Bowl Challenge (stage 1)[4], along with corresponding ground-truth expert manual annotations.

For consistency across cell-types and microscope technologies, we selected a subset of the 2019 Data Science Bowl Challenge dataset with dark backgrounds, resulting in 546 nuclei images, with a 80/20 train/validation split. *In situ* cell images are retrieved from the TNBC breast cancer MIBI (Multiplexed Ion Beam Imaging) dataset [9] for testing. Following suggestions from Keren et al. [9], we overlaid three nuclei marker channels (dsDNA, H3k27me3 and H3K9a) to generate each individual image. To evaluate our performance, we include the only two images (from Patient 1 & 2) which had manually annotated ground-truth masks [9]. For each 2048 × 2048 raw input images, we split each into 16 non-overlapping 512 × 512 patches, and further resize each patch into 256 × 256 to match our training image size. In summary, we trained on 437 (80% of nuclei images) *in vitro* nuclei images and evaluated our method on 32 *in situ* MIBI test images.

### 3.2 Experiment

#### *In vitro* training

We resized each image to 256 × 256, and converted it to grayscale. We trained the model for 50 epochs with early-stopping (patience count = 30) guided by the validation metrics. By default the learning rate is set as 1*e*^*−*3^. After training the model, we employed our multi-level watershed post-processing to achieve the final cell segmentation results.

#### *In situ* predictions

To evaluate our pipeline’s generalizability to *in situ* tissue data, we benchmarked it on the MIBI dataset. Table 1 compares the performance of our proposed method FPN+SAW+Multi-level Watershed using the manually annotated ground-truth masks, against U-Net, U-Net + SAW, and FPN + SAW, with or without Multi-level Watershed.

**Table 1:** Performance comparison on MIBI *in vivo* dataset.

| Model | Accuracy | F1 | IoU | AUC | Hausdorff |
| --- | --- | --- | --- | --- | --- |
| U-Net | 0.7494 | 0.7115 | 0.3091 | 0.6478 | 9.6090 |
| U-Net + SAW | 0.7543 | 0.7157 | 0.3168 | 0.6516 | 9.5787 |
| FPN + SAW | 0.7589 | 0.7244 | 0.3324 | 0.6606 | 9.4976 |
| U-Net + Multi-level Watershed | 0.8216 | 0.8114 | 0.5260 | 0.7624 | 8.6590 |
| U-Net + SAW + Multi-level Watershed | 0.8396 | 0.8338 | 0.5814 | 0.7934 | 8.4070 |
| **FPN + SAW + Multi-level Watershed** | **0.8435\*** | **0.8397\*** | **0.6021\*** | **0.8055\*** | **8.2529\*** |

\* $p < 0.05$ (paired t-test comparing to U-Net)

Across our four evaluation metrics, FPN + SAW outperforms base U-Net and U-Net + SAW on the test *in situ* dataset. Our proposed method (FPN + SAW + Multi-level Watershed), not only significantly improves the segmentation pixel-wise accuracy by 10% when compared to base U-Net, but also outperforms the other models that leverage the same post-processing technique.

Figure 2 shows the result predictions^5^ for patients 1,2 where our method is able to segments cells even in densely populated regions without including any images from this tissue type or technology in the training set. Figure 3 shows a zoomed-in region in patient 1 illustrating that our pipeline can successfully separate most of adjacent cells, some of which are difficult to separate even visually from the raw images (i.e. red box regions). In addition to the quantitative results from Table 1, this image highlights the generalizability of our model from *in vitro* to *in situ* cells as it is able to use the geometric features learned from the *in vivo* training data to precisely predict accurate segmentation masks for the *in situ* test set.

**Figure 2:**
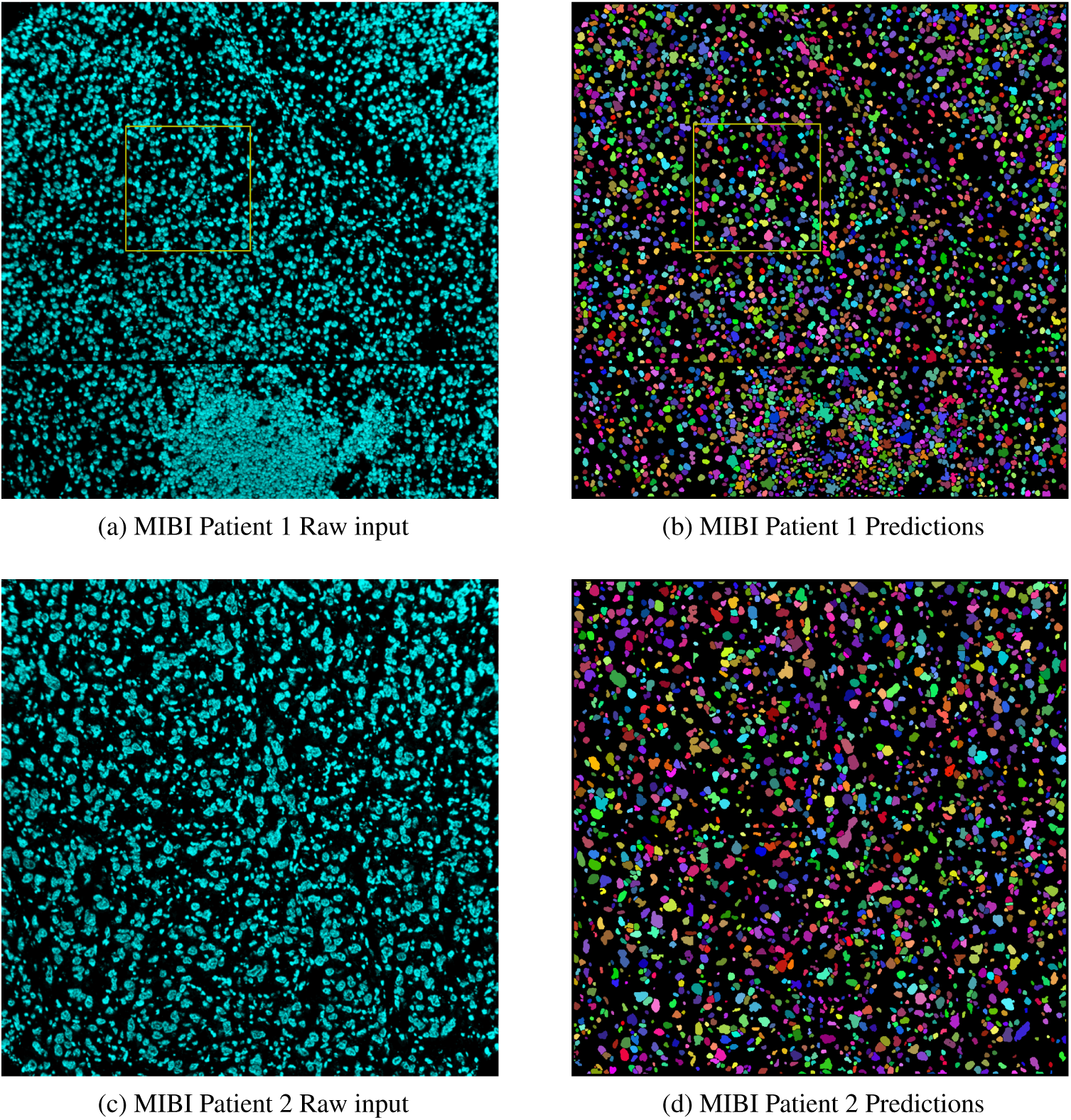
Panoptic predictions (right column) for patients 1,2 compared to actual image (left column) from MIBI dataset [9]. Yellow box indicates region zoomed-in in Figure 3.

**Figure 3:**
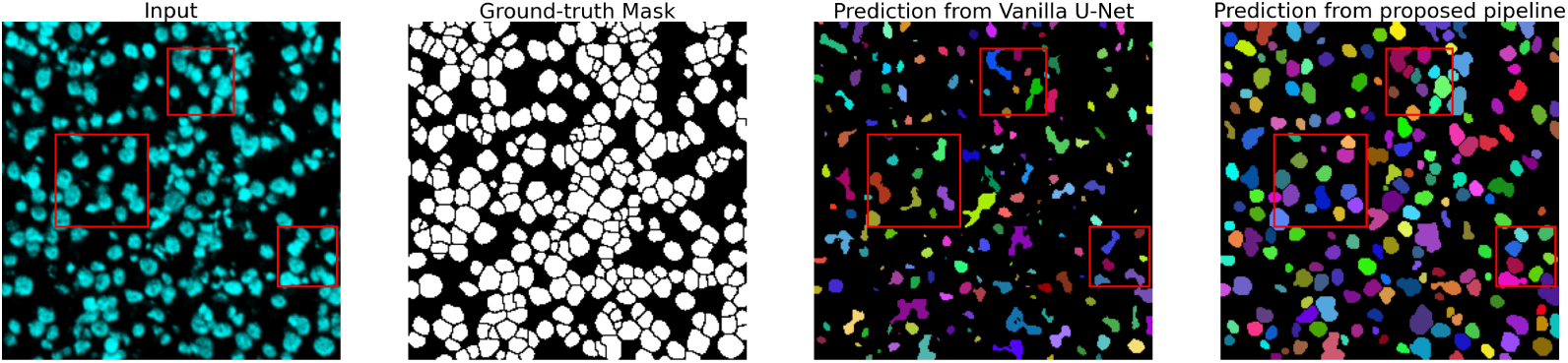
Zoom-in view of segmentation result for patient 1 in MIBI *in situ* dataset compared to ground truth annotations and prediction from base (vanilla) U-Net. Red boxes indicate regions with attached cells that are succesfully separated using our pipeline.

## 4 Conclusion & Future Directions

Our novel cell segmentation pipeline leverages shape-aware map pre-possessing and weights, and multi-level watershed post-processing to *in situ* tissue images captured with MIBI technology. To the best of our knowledge, this is the first method that can accurately segment *in situ* tissue data with pre-training solely on *in vitro* nuclei images, and this was accomplished with incorporating geometric features of cells into a deep-learning framework. Based on these promising results, our future directions will aim at expanding this idea by incorporating topological priors into the loss function [8] in addition to the SAW geometric features, leveraging the concept of persistent homology. Additional work down the line would be to extend our method to 3D segmentation and further test its robustness on single cell proteomics data as it becomes available.

## Footnotes

3 By definition, solidity of any close-loop shape *g* is the ratio of its area divided by its convex hull’s area: *Solidity*(*g*) = *Area*(*g*)*/Area*(*ConvexHull*(*g*))

4 Define *InverseDistanceTransform* as the operation of 0 - Euclidean distance transform from each pixel to its closest background. This generates a topography where the cell’s central regions have lower values than cell boundaries.

5 We applied the mask_rgb function in Cellpose [14] to generate colored masks

